# Application of a convolutional neural network to the quality control of MRI defacing

**DOI:** 10.1101/2021.06.24.449613

**Authors:** Daniel J. Delbarre, Luis Santos, Habib Ganjgahi, Neil Horner, Aaron McCoy, Henrik Westerberg, Dieter A. Häring, Thomas E. Nichols, Ann-Marie Mallon

## Abstract

Large-scale neuroimaging datasets present unique challenges for automated processing pipelines. Motivated by a large clinical trials dataset of Multiple Sclerosis with over 235,000 MRI scans, we consider the challenge of defacing - anonymisation to remove identifying facial features. The defacing process must undergo quality control (QC) checks to ensure that the facial features have been adequately anonymised and that the brain tissue is left completely intact. Visual QC checks – particularly on a project of this scale – are time-consuming and can cause delays in preparing data. We have developed a convolutional neural network (CNN) that can assist with the QC of the application of MRI defacing; our CNN is able to distinguish between scans that are correctly defaced, and can also classify defacing failures into three sub-types to facilitate parameter tuning during remedial re-defacing. Since integrating the CNN into our anonymisation pipeline, over 75,000 scans have been processed. Strict thresholds have been applied so that ambiguous classifications are referred for visual QC checks, however all scans still undergo a time-efficient verification check before being marked as passed. After applying the thresholds, our network is 92% accurate and is able to classify nearly half of the scans without the need for protracted manual checks. Even with the introduction of the verification checks, incorporation of the CNN has reduced the time spent undertaking QC checks by 42% during initial defacing, and by 35% overall. With the help of the CNN, we have been able to successfully deface 96% of the scans in the project whilst maintaining high QC standards. In a similarly sized new project of this scale, we would expect the model to reduce the time spent on manual QC checks by 125 hours. Our approach is applicable to other projects with the potential to greatly improve the efficiency of imaging anonymisation pipelines.

## 1. Introduction

A collaboration between Novartis and the University of Oxford’s Big Data Institute (BDI) has been established to improve drug development and healthcare through the application of artificial intelligence and advanced analytics. Large, multidimensional datasets – consisting of clinical, imaging and omics data – are integrated and analysed to improve patient prognoses and identify early predictors of disease. A dedicated research informatics framework has been developed that allows this data to be captured, anonymised, explored and integrated into databases (Mallon et al., 2021). Currently, the collaboration focuses on two main therapeutic areas: Multiple Sclerosis (MS), and autoimmune diseases treated with the interleukin (IL)-17A antibody secukinumab (IL17). Data from over 50,000 patients is available across both projects, and the MS project will utilise both clinical and brain magnetic resonance imaging (MRI) data – a key part of the MS project, with over 235,000 scans from over 12,000 unique subjects. Many of the subjects have longitudinal MRI data over several years and corresponding clinical information, enabling the progression of disease to be studied (Dahlke et al., 2021).

Before the MRI data can be used for research purposes it is necessary to homogenise the data to a single format, and to anonymise all identifying patient data, including metadata and any facial features that are present in the image data. While MRI metadata (stored within DICOM tags in raw MRI data) can be readily anonymised, through deletion or modification of selected tags, the anonymisation of the image data itself is more complex. Two commonly used approaches are skull stripping and defacing. Skull stripping involves removing all non-brain tissue from a scan, and can be implemented using a number of methods (e.g. Ségonne et al., 2004; Shattuck et al., 2001; Smith, 2002). However, skull-stripping methods often require considerable fine-tuning when applied to large datasets containing scans of variable image quality (Fennema-Notestine et al., 2006), and additionally, skull-stripped images can sometimes be unsuitable when similar algorithms are required for processing images downstream of de-identification work (Schimke et al., 2011). Defacing techniques retain non-brain tissue and can be implemented through shearing off parts of the face (e.g. Schimke and Hale, 2011), blurring the outer surface of the face (Milchenko and Marcus, 2013) and selectively removing areas of the face that contain identifiable facial features (e.g. Bischoff-Grethe et al., 2007; Alfaro-Almagro et al., 2018; Gulban et al., 2019).

Quality control (QC) processes are commonly employed when working with MRI data to ensure that the data is suitable for downstream analysis, and that artefacts in the data will not introduce bias into analyses (Reuter et al., 2015). Furthermore, the quality of defacing may also need to be QC checked (e.g. Marcus et al., 2013) to ensure that not only are any identifying facial features correctly removed from the scan, but also to ensure that the brain anatomy has not been damaged. The choice of defacing software is important as the successful application can be variable, particularly when the data has been acquired at different sites, with different acquisition parameters (Theyers et al., 2021). Additionally, the choice of software can impact the results of downstream analyses (Bhalerao et al., 2021; Schwarz et al., 2021). If high standards of QC checks are not employed, then there is potential risk of patients being identified through photographic visual comparisons (Prior et al., 2009), facial recognition software (Mazura et al., 2012; Schwarz et al., 2019, 2021), and the facial reconstruction of inadequately defaced MRI scans (Abramian and Eklund, 2019; Schwarz et al., 2021). Many defacing methods have been developed on high resolution, research quality scans. In this collaboration, which utilises MRI scans from global clinical trials, the scans are typically lower resolution, were captured from a large number of sites, and due to the longitudinal nature of the data, some scans were acquired over 15 years ago. Therefore, due to the potentially high levels of variation in the quality of the data, there is greater potential for variation in the successful application of defacing to these scans. As a consequence of this, thorough QC checks are necessary to ensure data is correctly anonymised.

While visual QC checks are commonly employed to assess the quality of MRI scans, when undertaking projects containing tens of thousands, or in the case of this project hundreds of thousands of scans, these manual checks become impractical. The time-consuming nature of the checks can cause considerable delays between receiving data and having the data research-ready. Furthermore, those undertaking the QC may become fatigued and more likely to make errors. Citizen scientists can be used to assist with visually QC checking MRI data (e.g. Keshavan et al., 2019), but this is not suitable when anonymisation checks are being undertaken. Automated methods have been developed to assist with the QC of MRI data, utilising methods including univariate classifiers (Mortamet et al., 2009), support vector machines (SVMs; Pizarro et al., 2016), random forests (Klapwijk et al., 2019), neural networks (Litjens et al., 2017), or a combination of classifiers (Alfaro-Almagro et al., 2018; Esteban et al., 2017). Automated QC methods may not perform adequately on all data - meaning that it may still be necessary to conduct some manual QC checks. However, allowing automated methods to handle the clear cases allows manual QC to be focused on data that is more difficult to classify. Additionally, automated methods must be generalisable so that they can process heterogeneous data. Despite the prevalence of automated models used to QC MRI data, these are typically focused on QC checking image quality or analysis output, and currently no models exist for assisting with the QC of the application of MRI defacing.

In this work, we developed a convolutional neural network (CNN) to assist with the QC checking of defaced MRI images from a large dataset containing images from numerous sources and of variable quality. Our network was developed using 24,000 pre-classified renders of defaced scans. Following development of the CNN, we evaluate model performance to select strict probability thresholds that convey high levels of accuracy, allowing for reliable automatic classification and for manual QC checks to be targeted towards problematic scans. We also describe the integration of the CNN into a pre-existing image anonymisation pipeline, complementing our existing manual QC processes, and including the adoption of time-efficient verification checks to protect patient anonymity. Furthermore, we evaluate the implications of implementing the CNN with regard to time saved compared to fully manual QC checks, and discuss the potential for the adoption of machine learning approaches to improve efficiency of MRI anonymisation pipelines.

## 2. Material and methods

### 2.1. MRI data

MRI data was available from 24 Novartis clinical studies, with the entire dataset containing data from over 12,000 subjects and over 235,000 MRI scans. The majority of scan sessions contained T1-weighted (with and without Gadolinium contrast enhancement), T2-weighted and proton density modalities. Some subjects also had T1-weighted 3D acquisitions, diffusion-weighted, FLAIR, and magnetisation transfer scans. MRI scans were of variable quality as they were captured from over 1,000 unique MRI scanners, with some scans captured over 15 years prior to the start of the collaboration. The majority of scans had a 256 × 256 or 192 × 256 acquisition matrix, with 46 or 60 slices. Prior to anonymisation, facial features are readily discernible in these scans.

### 2.2. MRI defacing pipeline prior to CNN development

MRI data was initially transferred to a dedicated, secure anonymisation environment for defacing and the anonymisation of any remaining confidential patient data. The anonymisation environment was only accessible to a very small team of dedicated scientists not otherwise involved in the research. A bespoke pipeline (Fig. 1) was built to handle the processing of this data into a consistent format, ready for downstream research. Data was initially converted to the Neuroimaging Informatics Technology Initiative (NIfTI) format using the DICOM conversion software HeuDiConv (Halchenko et al., 2019), and structured using the Brain Imaging Data Structure standard (BIDS; Gorgolewski et al., 2016) - a standard for brain MRI datasets, that is widely used within the neuroimaging research community. Anonymised metadata was preserved in a JSON file that accompanied each NIfTI file.

**Figure 1:**
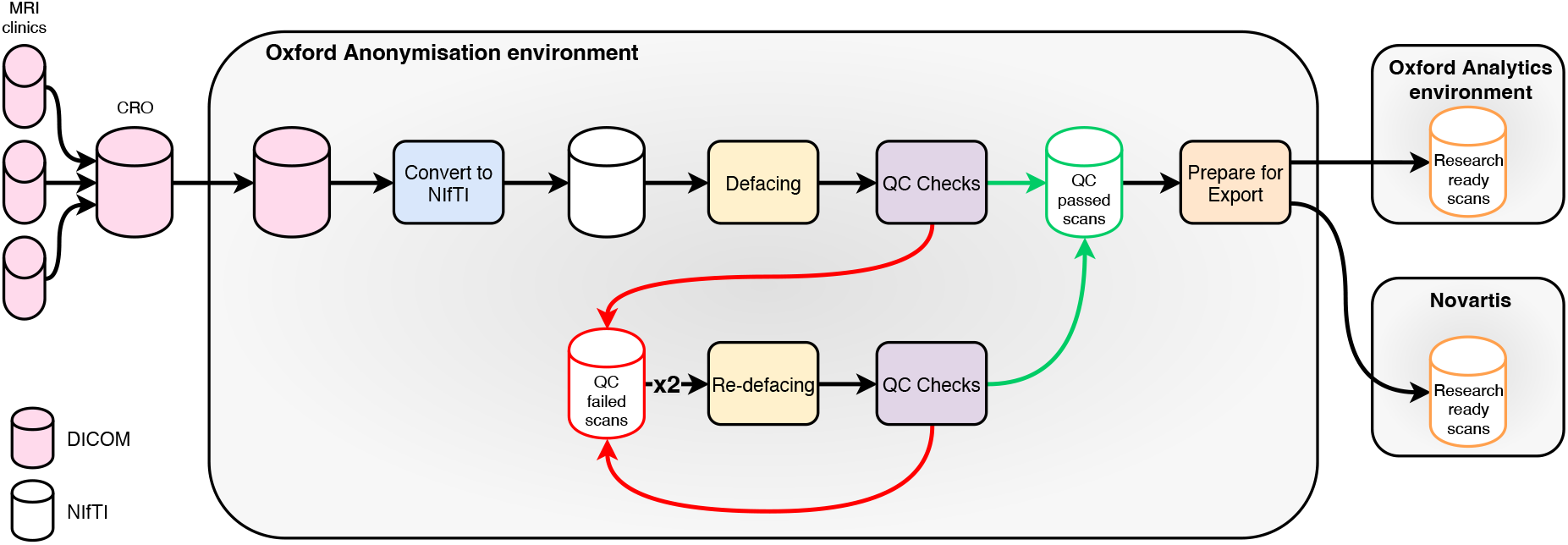
Flowchart showing the anonymisation pipeline for MRI data.

During the conversion process, a select number of DICOM tags are extracted and added to the JSON files by the converter; these tags provide fields that may be necessary for downstream analysis. These tags are selected by the conversion software and include tags recommended for inclusion by the BIDS specification. Some tags that were included by HeuDiConv were completely removed from all JSON files; this included some free-text fields (e.g. ‘Image Comments’) and details that could identify institutions. Unique values for JSON tags were reviewed to ensure that no un-anonymised data had inadvertently been included in the JSON files.

After conversion to NIfTI format, each scan was defaced. During this process, identifying facial features, specifically the ears, eyes, nose and mouth, are removed from each scan. Defacing was implemented using the fsl deface software (Alfaro-Almagro et al., 2018), which is known to perform well at preventing re-identification and minimise interference with downstream analysis methods when compared to other defacing software (Schwarz et al., 2021). Unlike many of the other available defacing methods, fsl deface removes the ears as well as other facial features. Following being defaced, all scans were QC checked to ensure that the defacing had been applied correctly. Scans were classified as one of four categories: pass (scan defaced correctly), deep (defacing went too deep and encroached on the brain), shallow (defacing did not go deep enough and facial features were still visible), failure (broad category for complete registration failures and scans containing unfixable errors; defacing failures that do not fit into the ‘deep’ and ‘shallow’ categories). Prior to the development of the neural network, all QC checks were carried out manually, using a HTML page generated by a Python script. Each of the scans were visualised as two PNG images of 3D renders of the scan (left and right oblique views) which were generated using the fsl gen 3D script (Alfaro-Almagro et al., 2018). Using the HTML page, each scan was classified by clicking the button that corresponds to one of the above categories (Fig. 2).

**Figure 2:**
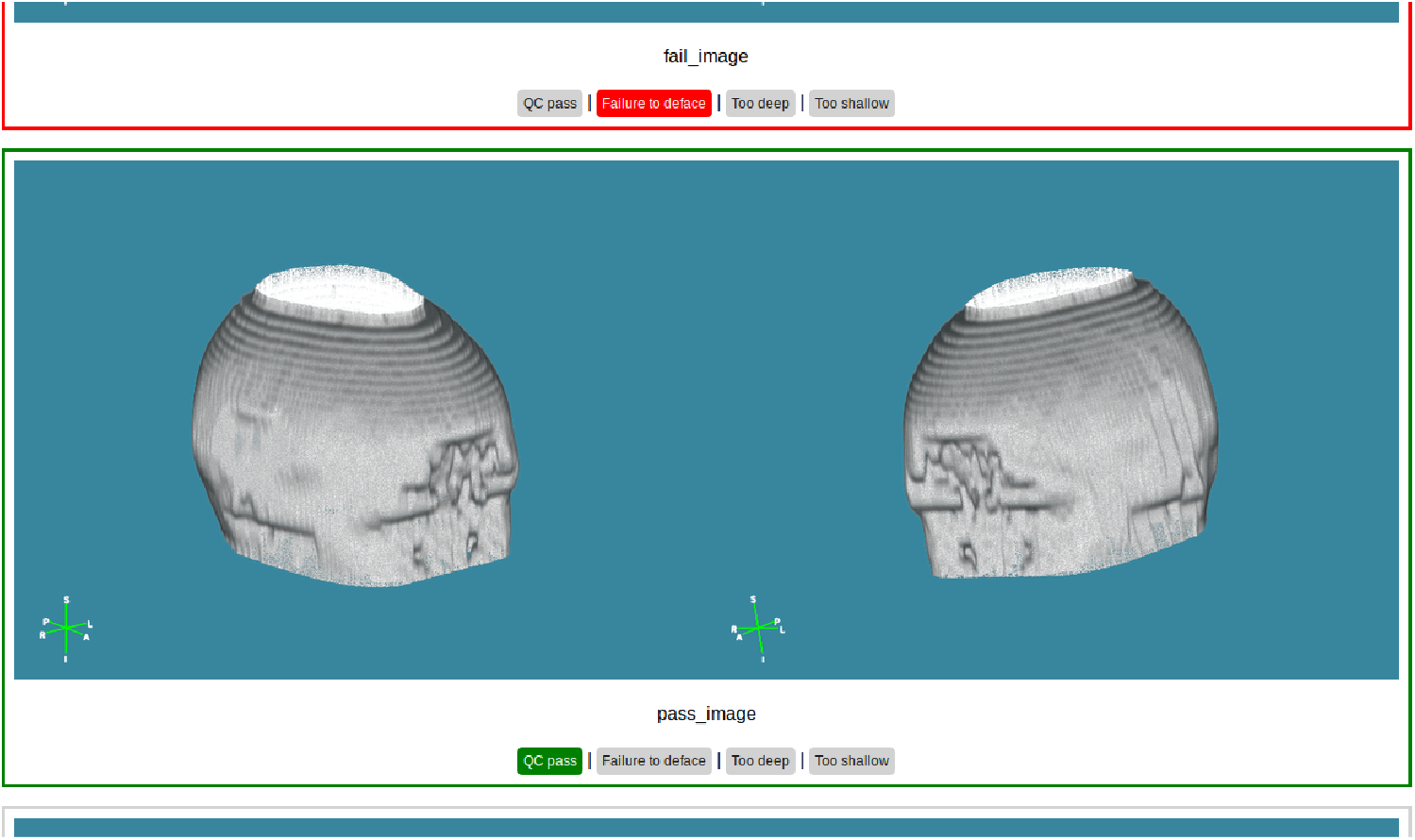
Usage of the html page when undertaking manual QC checks. Note that the renders in this figure are generated from a publicly-available sample scan from I Do Imaging (2017).

After the initial QC checks were completed, remedial defacing was undertaken on scans which did not pass, a process that we called re-defacing, where custom parameters were selected by considering the type of scan and the previous QC classification(s). When defacing was classified as too deep or too shallow, the defacing mask was nudged anteriorly or posteriorly respectively during the re-defacing. The application of bias correction and/or reducing the fractional intensity was used when a defacing attempt had been classified in the ‘failure’ category. The re-defacing was also applied in an additive manner. For example, if a scan had been classified as ‘shallow’ during initial QC checks, and was still classified as ‘shallow’ after the first round of re-defacing, then the mask would be nudged even more posteriorly during the second round of re-defacing. Following each round of the re-defacing, scans were once again QC checked using the same protocol as with the initial defacing. Up to two rounds of re-defacing were undertaken to ensure that as many scans as possible passed the defacing stage. On average, after initial defacing 69% of scans had passed QC checks, but 96% of scans were successfully defaced after two rounds of re-defacing. All anonymised NIfTI scans, and accompanying JSON metadata files, that passed the QC checks were then prepared for export to a separate, analysis-focused environment at the BDI and also back to Novartis, ready to be included in downstream research and analysis.

### 2.3. CNN: Data selection and pre-processing

As over 100,000 scans had been put through our anonymisation pipeline and manually QC checked prior to the development of the neural network, a large quantity of labelled scan data was available for use in developing the CNN. However, not all of this data could be used in network development for two reasons. Firstly, as the majority of prior manual QC classifications were for scans classified as ‘pass’ (75%) and to a lesser extent ‘shallow’ (15%), there was considerably less ‘deep’ (6%) and ‘failure’ (4%) classification data, so in order to keep the proportion of data from each class balanced (at least initially; see below) a smaller subset of the available scans were used. Secondly, as most subjects usually have multiple scans (usually 8+, but up to 60) and some of these scans had been re-defaced and therefore QC checked multiple times, it was preferable not to use all available scans to reduce the chance of the CNN overfitting to the anatomy of subjects appearing in the dataset multiple times.

Initially, 16,000 scans were used to develop the model, with 4,000 scans per class, which were split 60:20:20 between training, validation and test sets. Due to the initial models performing better on the ‘deep’ and ‘failure’ classes, the proportion of ‘shallow’ and ‘pass’ scans in the dataset was increased to 8,000 scans per class, giving 24,000 scans in total. Scans for each class were randomly selected from those available, and included scans from multiple clinical studies, all available scan modalities, and they were of varying image quality, representing the variability in the entire dataset.

A single 2D image was used as the input for the CNN. Each image was a horizontal concatenation of the two renders used in the manual QC checks, which were cropped to remove parts of the background that did not contain parts of the anatomy in any scans. The rationale for using the 2D renders over the 3D scans was as follows: the average NIfTI scan in this project contains over 3 million voxels, most of which is not data that is applicable to assessing the quality of the defacing and would be computationally expensive to process in its entirety. Training needed to be computationally efficient as GPUs were not available within the anonymisation environment, and it was not possible to move the data externally for this purpose. In addition, while image registration could be used to extract the head/face region (and downsample the 3D data), defacing failures are often the result of poor image registration. Therefore, by using 2D images of the renders, visual information that is pertinent to the effectiveness of the defacing is retained in a more compact format, and the style of the renders is consistent regardless of image quality. The renders were input into the CNN as RGB data for the following reasons. Firstly, the renders are generated with a coloured background for the manual QC checks; not having to convert these renders to greyscale streamlines the integration of the CNN into the defacing pipeline. Secondly, it is possible that the CNN could use the coloured background to detect edges of the render, including cases where the defacing mask has been applied incorrectly leaving large holes in the scan. Thirdly, the impact of using three channels was minimal on performance, with only a moderate increase in training time (∼10%).

### 2.4. CNN: Network development

The CNN was developed using the ‘keras’ library (Allaire and Chollet, 2018) for R (R Core Team, 2018). Images were input to the network as 165 × 270 × 3 (height x width x depth) tensors with values scaled between 0-1. The image order was shuffled and image augmentation (zoom, horizontal flipping) was used on the training data set to reduce overfitting; augmentation was not applied to validation or test data. The base of the network (Fig. 3) contained four convolutional layers, each using a rectified linear unit (ReLU) activation function and a kernel size of 3 × 3. Each convolutional layer was followed by a max pooling layer to downsample the data. After flattening, two dropout and three densely connected layers were interspersed. Both dropout layers had a dropout rate of 0.2. The first two dense layers used ReLU activation, with the final dense layer using softmax activation with an output size of four – one for each of the four classes. The network was optimised using a root mean square optimiser (RMS) with the learning rate set to 0.00008, and categorical cross-entropy was used as the loss function. The network was trained for 100 epochs, with 576 steps and a batch size of 25, so that each epoch contained all 14,400 training images. During each epoch the model was run on the validation set, with 192 steps and a batch size of 25. A callback was used to save the best performing model during training, using minimised validation loss as the criterion.

**Figure 3:**
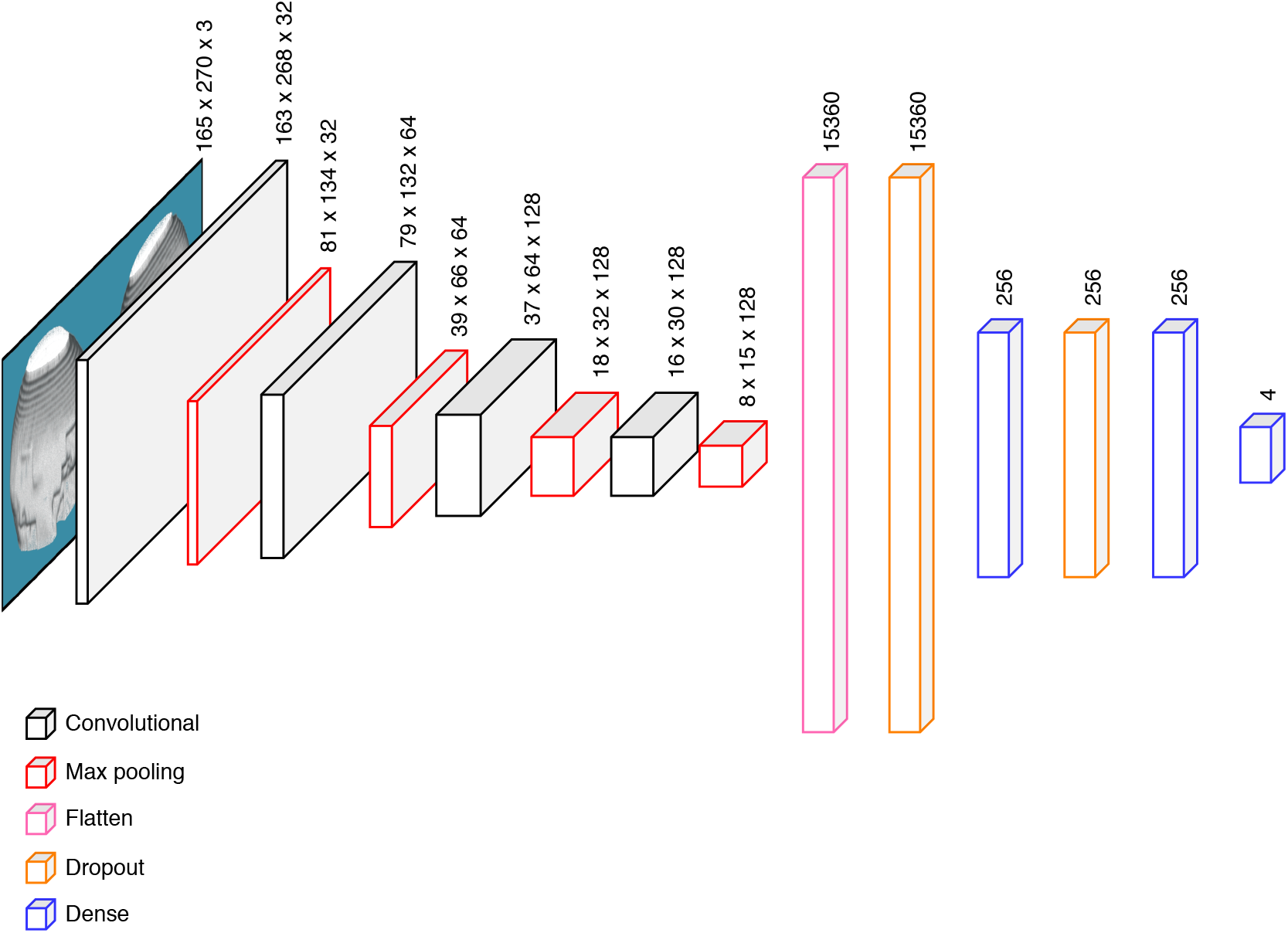
Network diagram of the CNN.

The assignment of classifications was visualised using Gradient-weighted Class Activation Mapping (Grad-CAM; Selvaraju et al., 2017). This approach allowed for the activation of a class for an input image to be visualised, which shows what features of the MRI renders the CNN is using to assign classifications. The Grad-CAM method was implemented in R using the approach of Chollet and Allaire (2018), where a semi-transparent heatmap of the predicted class activations from the final convolutional layer is superimposed on top of the input image. For illustrative purposes, a publicly-available sample T2w MRI dataset (I Do Imaging, 2017) was used for the images included here; the parameters of the defacing script were modified to produce an example of each of the four classes.

Multi-class receiver operating characteristic (ROC) curves, and the corresponding area under the curves (AUC) were calculated using the method of Hand and Till (2001), as implemented in the pROC R package (Robin et al., 2011).

### 2.5. Probability threshold selection

Before the network was incorporated into the defacing pipeline, cut-off probability thresholds were selected so that only classifications with high confidence would be accepted. The adoption of strict cut-off thresholds was necessary to ensure that inadequately defaced scans are not marked as ‘pass’, and that incorrectly defaced scans were allocated to the correct class to allow for the appropriate parameters to be applied during remedial re-defacing work. Performance across classes was evaluated to select thresholds that will deliver acceptable performance upon implementation into the defacing pipeline. Cut-off probability thresholds were evaluated using metrics including sensitivity, specificity, and volume of data that would be classified at different thresholds (between 0.25 – 1.00 at 0.05 increments) for each class on the test dataset. Classifications were assigned based on the class with the highest probability. During the implementation phase (see below), classifications which did not surpass the threshold were discarded and these scans would go through the manual QC checks instead. Therefore, the metrics used to assess the impact of applying different thresholds only include scans with accepted classifications. For example, if at threshold *x* classifications for 600 (out of 1,000) scans reached the threshold, the sensitivity and specificity would be calculated for only those 600 scans that would be classified, and not the entire set of 1,000 as we know that the remaining 400 scans will be manually QC checked.

### 2.6. Integrating the CNN into the defacing pipeline

The CNN was integrated into the defacing pipeline (Fig. 4), and was used at the initial defacing and the two re-defacing stages. Following the defacing of all scans in a study, a concatenated image of the two renders was generated to match the format of the images used in training the network. Classifications for the three non-pass categories were accepted when they exceeded the cut-off probability threshold for a class. Pass classifications that surpassed the pass cut-off threshold were preliminarily accepted, pending a visual verification check before being fully accepted. The visual verification check is a quicker version of the full manual QC check, where the images are displayed in a gallery view that can be swiftly browsed and only those images where the pass classification is not agreed with are flagged. Any scans that were flagged during the verification check, and the remainder of the scans that did not reach any of the cut-off thresholds, were then QC checked using the original manual method.

**Figure 4:**
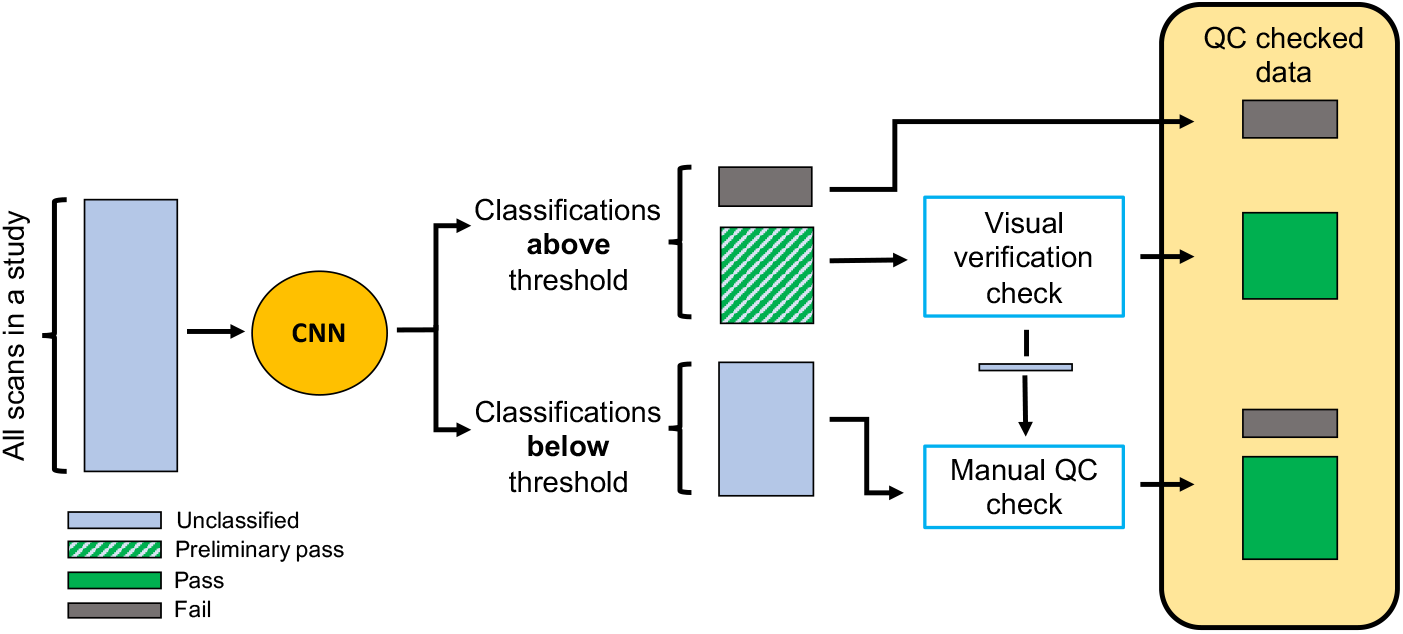
Flowchart showing how the CNN has been integrated into the anonymisation pipeline. This process takes places within the ‘QC checks’ part of the anonymisation pipeline (shown in figure 1).

After the CNN was incorporated into the pipeline, its impact was assessed relative to the baseline manual QC process that was used prior to development of the CNN. The efficiency savings by using the CNN, compared to fully manual QC checks, was determined by calculating the time savings for a hypothetical study containing 10,000 scans. The performance of the CNN (e.g. proportions of scans classified in the four classes, the proportion of CNN assigned passes accepted during verification checks) was based on the performance of the CNN on data processed by it. The time taken to perform the manual and verification checks was determined by calculating the average time it takes to check a single scan in a batch of 250 scans.

## 3. Results

### 3.1. Network performance

When the test set was evaluated using the CNN, accuracy of 0.76 with a 0.56 loss was achieved; this is similar to the values obtained during training on the validation set (0.77 accuracy and 0.56 loss). The model performs very well when distinguishing between deep and shallow classes (AUC = 0.97; Fig. 5a), and performs reasonably well distinguishing between failure and shallow classes (AUC = 0.90), and between deep and pass classifications (AUC = 0.87). The CNN performs poorest when distinguishing between failure and pass data (AUC = 0.77).

**Figure 5:**
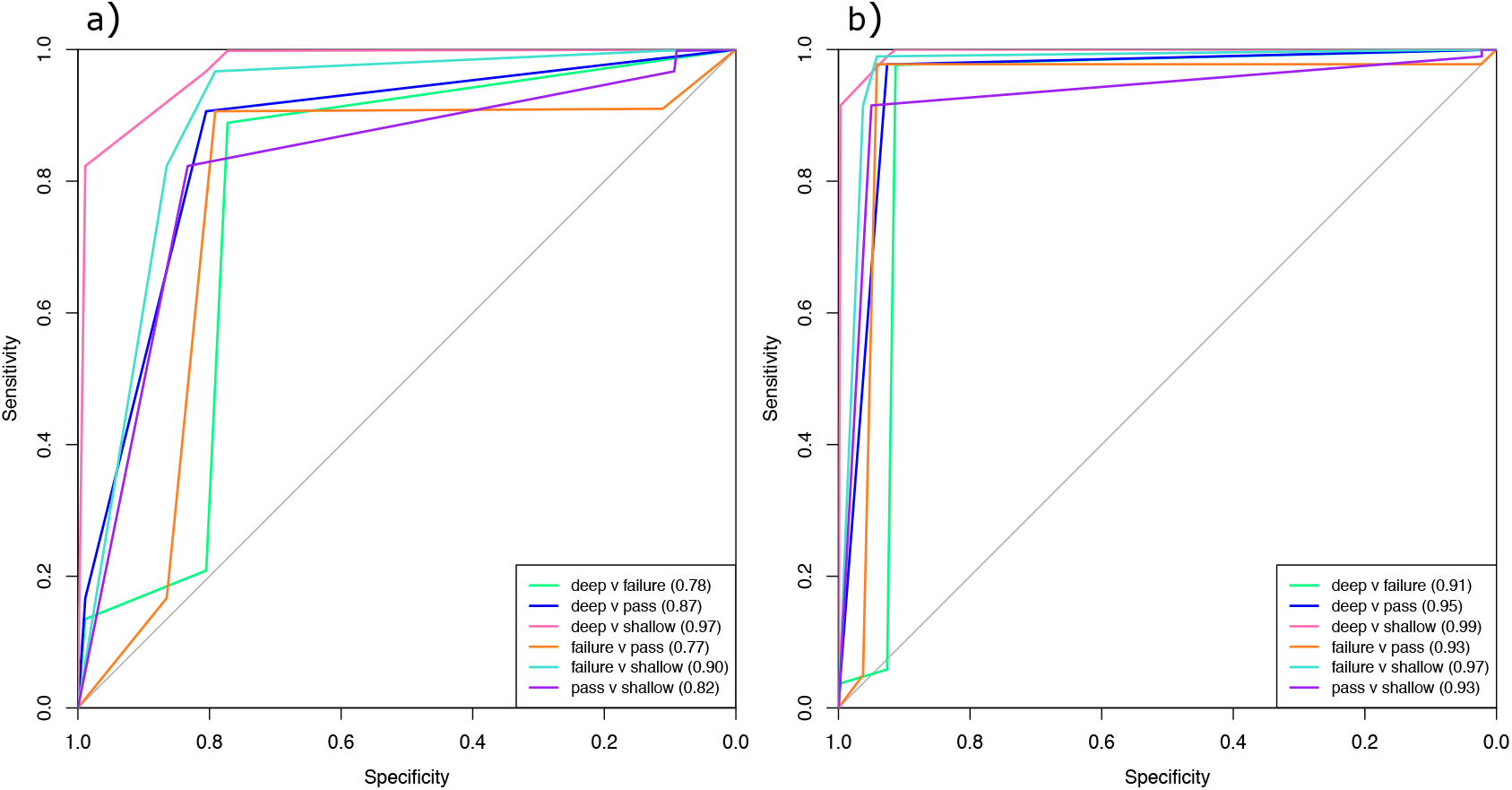
Multi-class ROC curves when a) all classifications are accepted, and b) when the thresholds are applied. Each curve represents one pair-wise comparison. The AUC is shown in the legend for each comparison.

The features that the CNN uses to assign the classes can be visualised using the Grad-CAM heatmaps (Fig. 6). For the ‘pass’ scan (Fig. 6a) activation is highest just below one of the eyes, but there are also high levels of activation above the forehead. With a complete defacing failure, where behind the face has been defaced and the defacing has gone quite deep into the sides of the head (Fig. 6b), there is very strong activation around a hole in the forehead. Other areas of the head where the defacing has not been adequately applied are also activated. In a scan where the defacing has gone very deep and intersected the brain (Fig. 6c), there is strong activation at the front of the brain and around the strong angular lines where the mask has cut through the sides of the front of the head. Interestingly, only one render shows any activation. The ‘shallow’ scan, in which the eyes are still visible (Fig. 6d), has very strong activation around the orbits, and high levels around the whole face. In particular the ‘L’-shaped cuts where the defacing mask has started to cut into the area of the brows, and below the eyes show strong activation.

**Figure 6:**
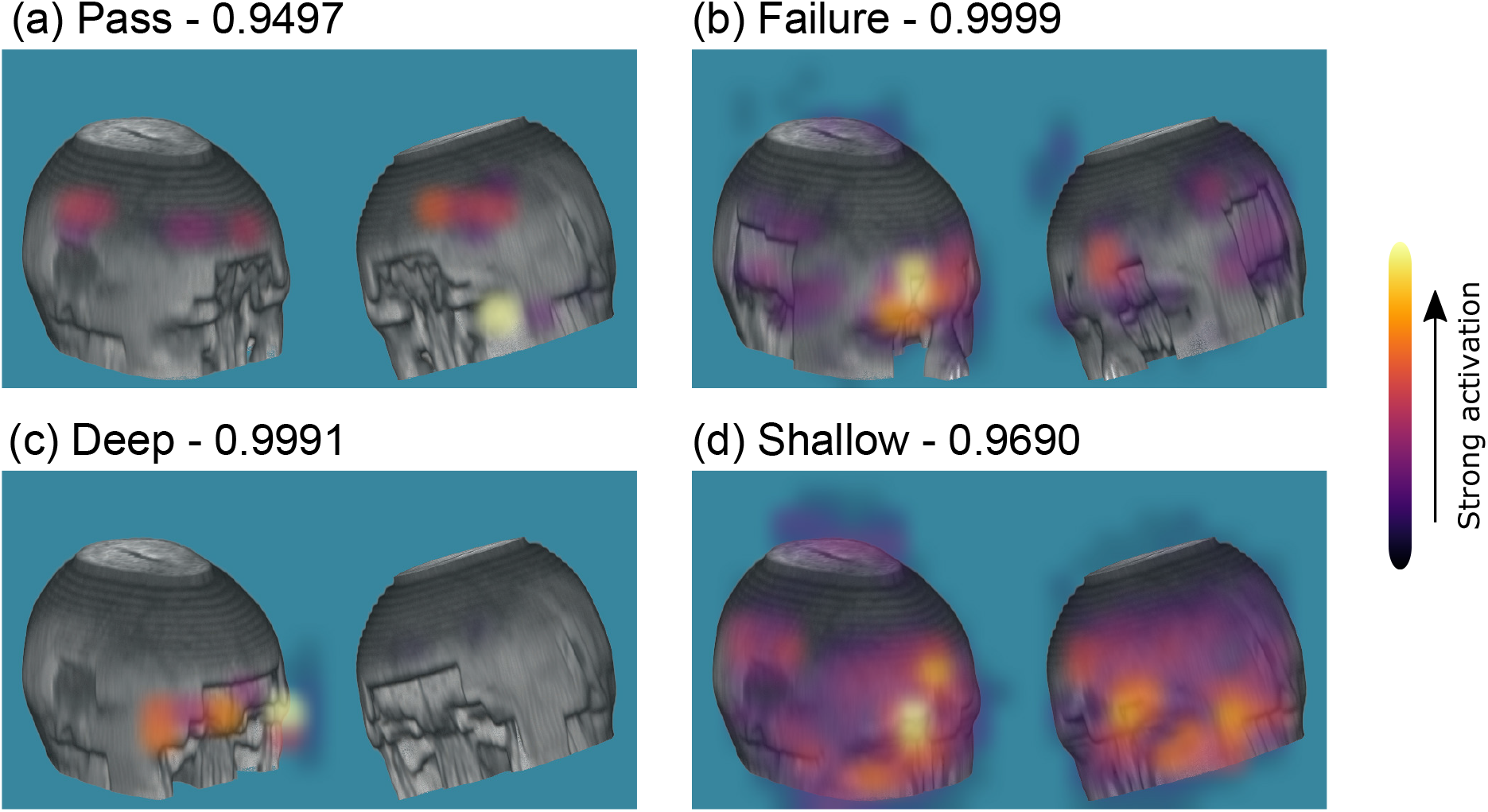
Grad-CAM heatmaps for representative scans of each of the four classifications, a) pass, b) registration failure, c) deep, and d) shallow. The CNN probability that each image belongs to the class is shown above each image. Each image was produced from a publicly-available sample scan from I Do Imaging (2017) with the defacing parameters modified to produce different outcomes.

The CNN is able to classify 1,000 images in 51 seconds (0.051 seconds per image), including the production of CSV files recording the classifications, and moving images to sub-directories based on assigned classes. When the number of images is increased to 10,000 the processing time decreases to 0.048 seconds per image (475 seconds in total).

### 3.2. Probability threshold selection

As the probability thresholds become less strict, the sensitivity and specificity decline in a similar pattern for the ‘pass’, ‘deep’ and ‘shallow’ classes although the specificity remains more consistent for the ‘deep’ class (Fig. 7). While there is a much greater decline in the sensitivity for the ‘failure’ class compared to the other three classes, the specificity remains very high (*>* 0.97) regardless of the threshold. When strict thresholds (0.95) are applied, the proportion of data which surpasses the selected thresholds is low for all classes (*<* 13%) except the ‘failure’ class where it is 46%. After examining the overall performance of the network, a global probability threshold of 0.8 was selected for all classes, with the exception of the ‘pass’ class. As scans assigned the ‘pass’ classification would undergo a visual verification check a slightly lower threshold of 0.75 was selected. Using multi-class ROC curves (Fig. 5b), the AUCs from all comparisons are ≥ 0.91, with most comparisons having AUCs ≥ 0.93. With the selected thresholds applied, 45% of data in the test set would be classified by the CNN, and 92% of these predictions match the labelled classes.

**Figure 7:**
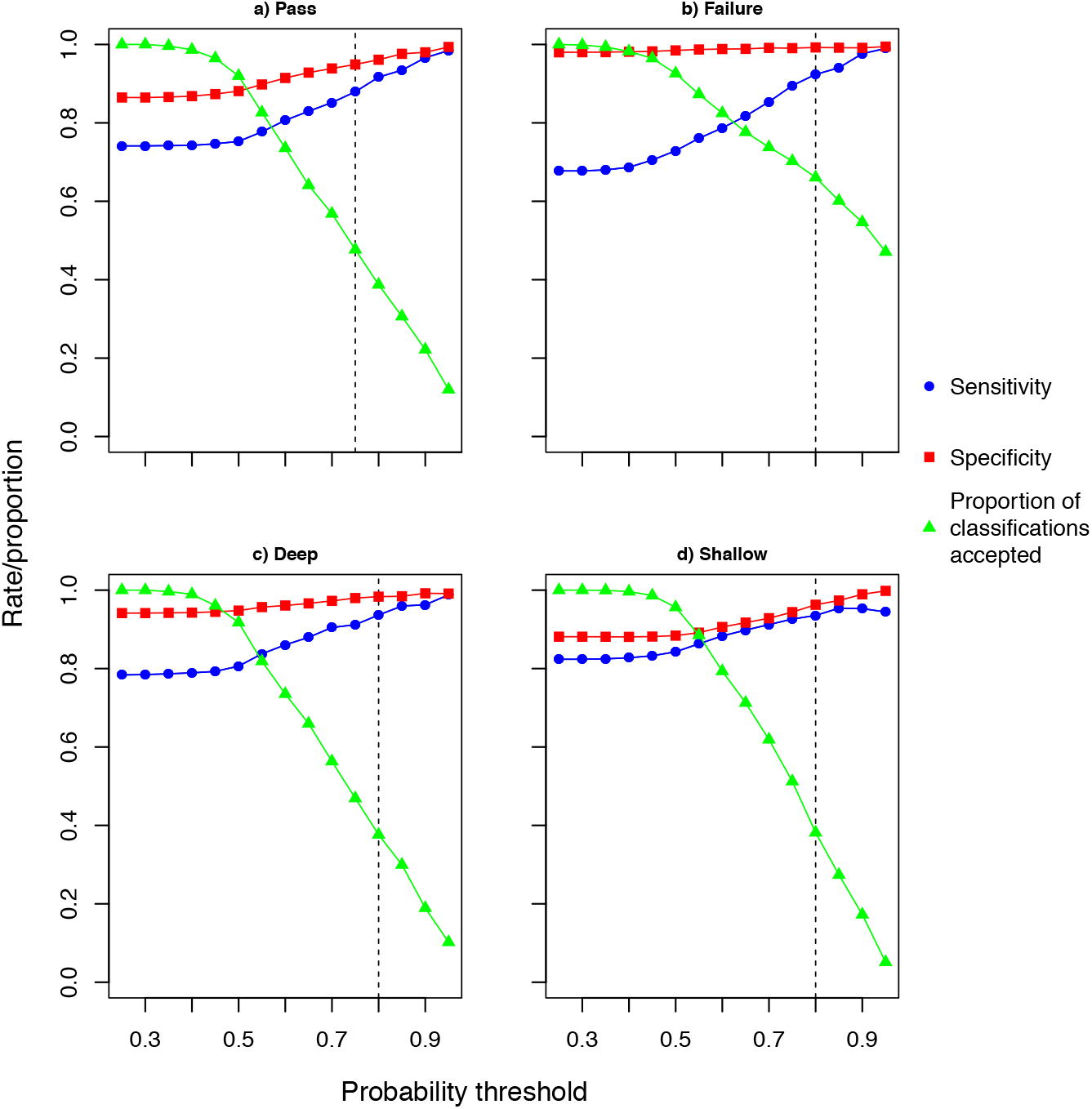
Plots showing how model performance on the test set varies when the probability threshold of accepted classifications is modified. Each class is shown separately: a) pass, b) registration failure, c) deep, and d) shallow. Sensitivity, specificity and the proportion of classifications that would be accepted are shown for each class (i.e. scans whose classifications would be discarded at each threshold are not included in the metrics). Vertical dashed lines show the thresholds that were used in the final model.

### 3.3. Integrating the CNN into the defacing pipeline

Since the CNN was integrated into the defacing pipeline, over 76,000 scans have been processed (Fig. 8), of which nearly 53% were from scans that had been through initial defacing, and the remaining 47% had been through re-defacing. The CNN classified 30% of scans as passed, 12% as shallow, 2% as deep, and 1% as registration failures; the remaining scans were flagged for manual QC checks as they did not surpass the applied probability thresholds. Of the scans assigned ‘pass’, 91% and 90% of these classifications were accepted following the visual verification checks in the initial and re-defacing stages respectively. Overall, 45% of the classifications generated by the CNN were accepted, however a larger proportion of the classifications (51%) were accepted for scans that were QC checked following initial defacing, and a much smaller proportion of classifications (39%) were accepted for scans that had been through re-defacing. Of the total proportion of scans classified by the CNN, 60% of scans were classified as pass during initial defacing, but this was much greater during re-defacing, where 78% of scans were classified as ‘pass’. This seems to be the result of a considerably smaller proportion of scans being assigned ‘shallow’ during re-defacing, due to the success of the revised defacing parameters that are applied during this stage.

**Figure 8:**
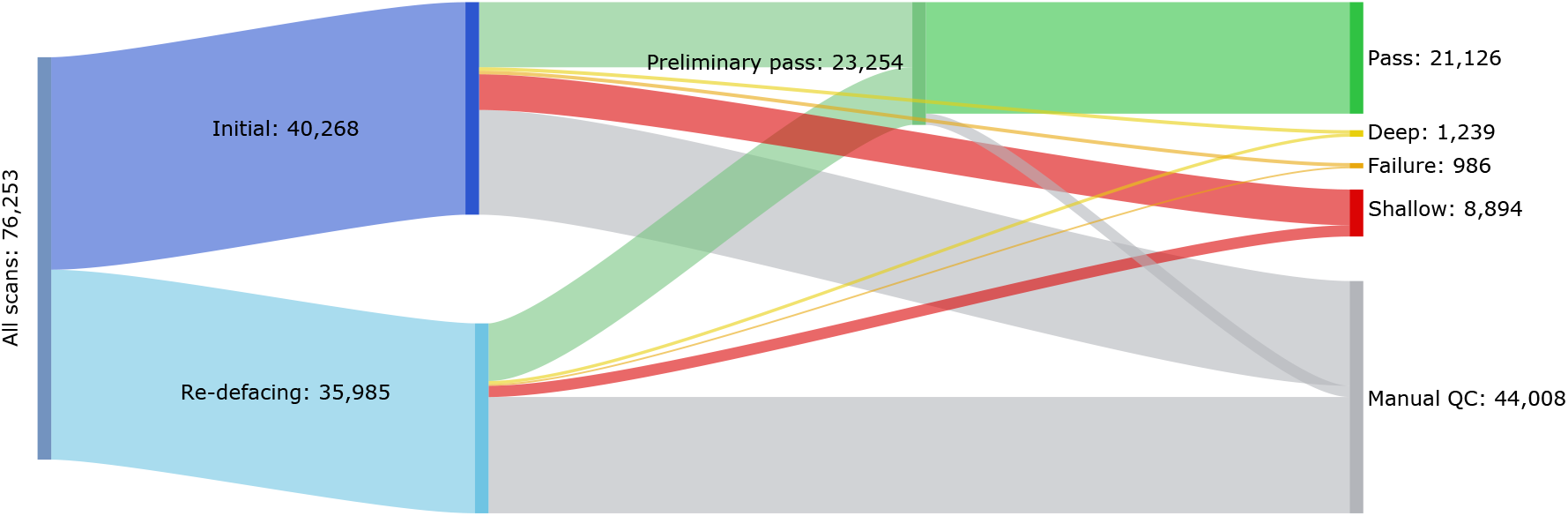
Sankey diagram showing scans that have been QC checked with the CNN incorporated into the anonymisation pipeline. Scans are split into those from initial defacing and re-defacing QC.

Incorporation of the CNN into the pipeline has allowed for a reduction in the amount of time spent on manual QC checks. Using summary data on scans that have been processed so far, it is possible to compute the time that is saved on an average study containing 10,000 scans (Fig. 9). Typically, 31% of scans in a study will need re-defacing, and 12% of the original scans will need a second round of re-defacing. On average, it takes a human scorer 3.8 seconds to visually QC check a scan, which includes time spent waiting for the image to load on the HTML page, looking at the image, and marking the classification. The visual verification checks are much quicker, taking 0.8 seconds per scan on average. Therefore, an average study with 10,000 scans would take 15.1 hours to manually QC check. With the addition of the CNN to the pipeline the time needed for manual and verification checks during initial defacing would be nearly halved to 6.1 hours (from 10.6 hours). During re-defacing, the time savings are less due to the CNN not performing as well on re-defaced scans (i.e. these scans are more commonly borderline between classes, or the subject’s anatomy or the scan quality makes defacing difficult). Verification and manual QC checks combined would take 2.5 hours during the first round and 1.2 hours during the second round of re-defacing when using the CNN. Prior to incorporation of the CNN, the manual checks would have taken 3.3 and 1.3 hours during the first and second rounds respectively.

**Figure 9:**
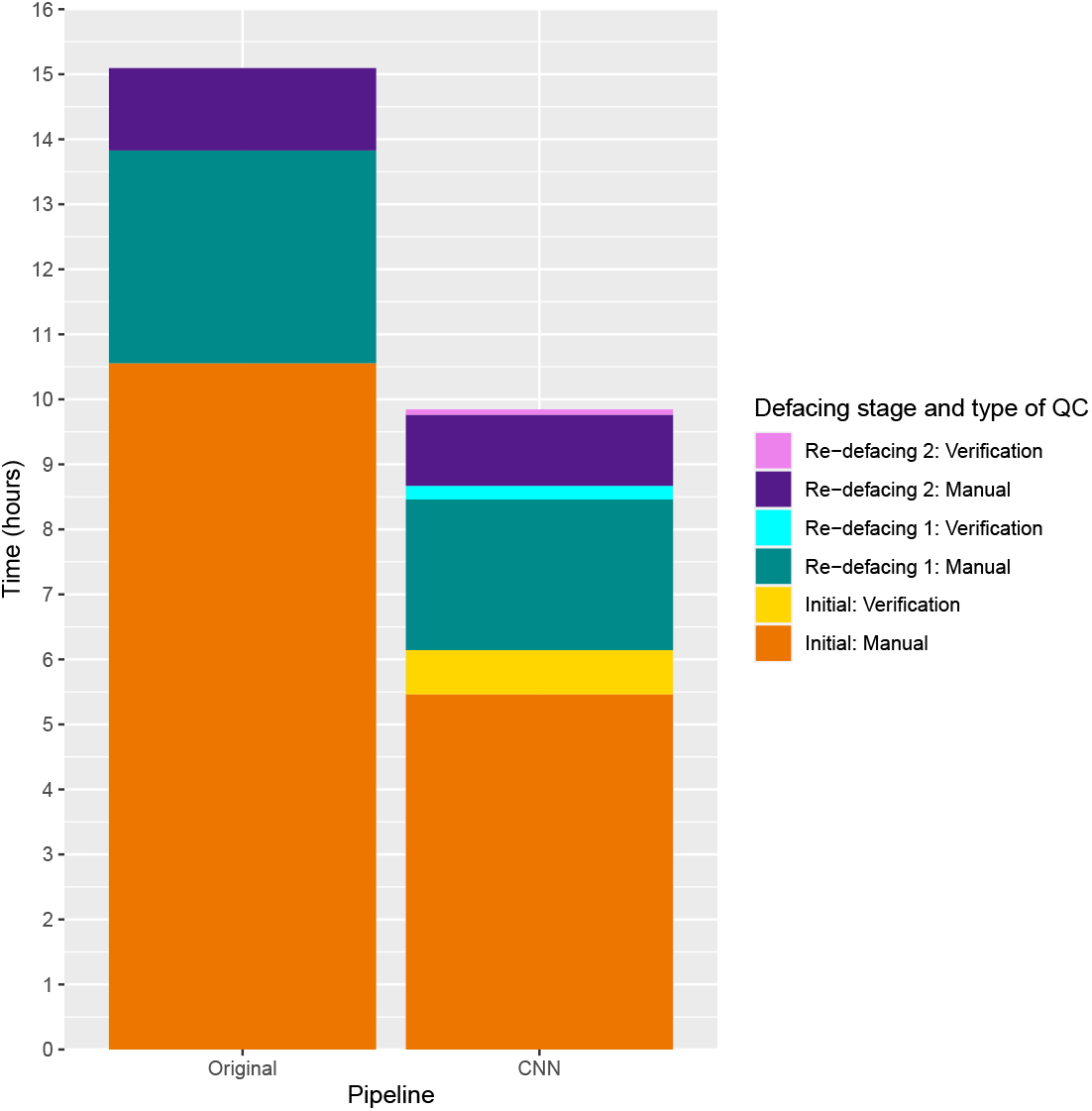
Stacked bar plot comparing the amount of time spent performing QC checks for an example study containing 10,000 scans, using the original fully manual pipeline and the pipeline with the CNN included.

In total, QC checks with the CNN incorporated into the pipeline would have taken 9.8 hours, instead of 15.1 hours prior to incorporation of the CNN into the pipeline, which is a 35% reduction in the time spent performing the checks. Additionally, with the CNN pipeline the entire QC process can be completed in less time than it takes to complete just the initial defacing QC using the original manual pipeline. During just the initial defacing - the most time-consuming part of QC checks - the time spent undertaking QC checks is reduced by 42% using the CNN pipeline. However, the benefits of the CNN are not as substantial during the re-defacing rounds where there is a 23% reduction during the first round, and only a 7% reduction during the second round. If the CNN pipeline was applied to a brand new project of the same size as this one (235,000 scans) the time spent undertaking QC checks could be reduced by approximately 125 hours.

## 4. Discussion

In this paper, we were able to develop a CNN that is able to perform QC checks on MRI defacing. Not only can the CNN identify scans that have been incorrectly defaced, it can also distinguish between the type of failure, which can assist when re-defacing these scans. We have shown how the CNN can be integrated within a defacing pipeline in a manner that delivers high standards of QC that protect the identity of subjects. Implementation of the CNN into the pipeline leads to a considerable reduction in the amount of time needed for manual QC checks, leading to faster data turnaround and an increase in capacity for MRI data anonymisation work.

Prior to applying probability thresholds to the final CNN, the network delivers test accuracy (0.76) which is similar to the performance of models applied to perform other forms of QC on large, multi-site MRI datasets (e.g. Pizarro et al., 2016; Esteban et al., 2017). However, as the CNN is used to QC check patient anonymity, higher levels of accuracy were required. The application of the strict probability thresholds conferred much higher test accuracy (0.92) but still allowed for nearly half of the MRI scans to be QC checked without the need for time-consuming manual checks. There is no existing comparable baseline method for conducting QC checks on the application of defacing. Recently, and since developing our own CNN, Bansal et al. (2021) have released a preprint documenting a binary classifier that can identify when defacing has or has not been applied to a scan. Similarly, our model can detect when defacing has been applied, but in addition our CNN can assess the quality of its application, allowing for the fine-tuning of parameters during re-defacing. This extra functionality was crucial for our QC processes and to allow us to successfully deface 96% of the scans that were part of this project. While this extra functionality is essential for our defacing pipeline, it presented some challenges. Classifying scans into the four discrete categories is not always straightforward, both for humans and the CNN, as scans often exhibit features characteristic of multiple classifications. For example, the front of the brain is often visible through the forehead in correctly defaced scans, or subjects with deeply-set eyes may show features of deep and shallow classifications. Furthermore, scans are often borderline between two categories (typically pass/deep or pass/shallow). In these cases, the CNN can be ambiguous with regard to assigning two (or more) classes. Taking all of this into account, there may be a ceiling for the accuracy of machine learning models that are used to assess the quality of defacing application, particularly when applied to datasets with variable image quality. While it may be possible to further improve the performance of the CNN, allowing for greater proportions of the data to be classified, it seems highly likely that manual QC checks will always be needed for scans that the CNN is unable to reliably classify.

Choosing appropriate thresholds was key to successful integration of the CNN into the defacing pipeline. The trade-off between volume of data processed and classification accuracy had to be considered with relation to the impact of the integration of the CNN into the pipeline. If less strict thresholds were applied, a greater volume of scans could be classified. However, even with the strict thresholds we applied here, up to 10% of assigned ‘pass’ classifications were not agreed with during verification checks. Further relaxing of the thresholds would most likely lead to a greater proportion of ‘pass’ assigned scans needing to be flagged for manual checks, increasing the average time to verify each scan, while making the verification checks less efficient and potentially eroding the time savings that the CNN provides. Furthermore, if the probability thresholds for the ‘deep’, ‘failure’, and ‘shallow’ categories were lowered then the quality of re-defacing attempts could be reduced. During the re-defacing, the custom parameters that are applied are highly dependent on the previous QC classifications. Inaccurate ‘failure’, ‘deep’ and ‘shallow’ classifications would lead to incorrectly applied re-defacing parameters, and a greater proportion of scans not passing the re-defacing, and consequently less data would be available for downstream analysis.

To protect patient identities, image anonymisation must remove all identifiable features as the presence of isolated facial features can compromise the anonymisation of image data. For example, a variety of machine learning approaches have been used to develop ear recognition tools, allowing ears to be used for biometric identification (Emeršič et al., 2017). With the application of visualisation methods like Grad-CAM we are able to show that our CNN has been trained to rely on the same anatomical features that human scorers use when deciding whether a scan has been defaced correctly. More precisely, we can verify that the network focuses on the eyes, ears, nose and to a lesser extent the mouth regions (defacing issues around the mouth region are less common). In particular, there was strong activation around the eyes for all classes, which is consistent with observations during the manual QC checks – that the eyes are one of the most common areas to exhibit problems. Also, there are rarely issues with the removal of the ears during the defacing, but it is reassuring that the CNN utilises features around this area of the head. Additionally, the CNN is able to correctly classify scans in the registration failure category despite the appearance of scans in this category being quite variable.

For projects like the Novartis-Oxford collaboration, that utilise hundreds of thousands of MRI scans acquired from sites around the world, one of the main challenges is being able to process the large volumes of data in a timely manner without compromising on the quality of the processing. While machine learning approaches, such as SVMs and CNNs, are regularly used when performing QC checks on MRI data, they are typically used to identify scans that are likely to be problematic when analysed (e.g. poor image quality, identification of artefacts). Our innovative approach of applying a CNN to assist with QC checking the anonymisation of image data highlights that there is potential for applying machine learning approaches to other stages of MRI processing pipelines. The aim of this paper was to develop a model that would assist with the QC of the defacing, and complement, but not replace visual QC checks, while maintaining high QC standards. This goal has been achieved with a reduction in the time spent undertaking manual checks by approximately 35% while maintaining the quality of the images that are passed through to the analytical pipelines. On a project of this scale the time savings are considerable and greatly improve the efficiency of the MRI anonymisation process. Therefore, other projects or platforms dealing with the challenges of anonymising large quantities of MRI data could also find that applying similar approaches as the one detailed in this paper can lead to substantial reductions in the time spent on manual QC processes. Unfortunately, sharing models used for QC checking anonymisation can be problematic when they are developed using patient data, due to the potential for un-anonymised data to be retained in these models (e.g. Fredrikson et al., 2015); it is necessary to restrict the storage and usage of these models to a secure environment. Additionally, different defacing methods will likely require bespoke models to capture the relevant features. However, if projects and platforms have existing QC classification data available, then impactful QC models - that are applicable to existing processing pipelines - can be developed even when using relatively simple deep learning architectures. Manual QC checks can be time-consuming bottlenecks, but with the application of approaches such as CNNs, this can be alleviated, expediting the availability of anonymised MRI data to researchers.

## Acknowledgements

The project, and the collaboration, was made possible through access to MRI data from Novartis’ MS clinical trials. We wish to thank Piet Arden, Frank Dahlke, and Karine Lheritier from Novartis for their assistance with providing access to data and supporting the collaboration. We wish to acknowledge the help of Mark Jenkinson in providing guidance with establishing the defacing pipeline; Stephen Gardiner, Ewan Straiton, and other members of the data wrangling team for their assistance in working with the MS MRI data; Anna Zalevski and Joanna Stoneham for project management; Adam Huffman, Geoffrey Ferrari, and Robert Esnouf for their work on the IT infrastructure and data transfers for the project.

## Ethics statement

Data used by the collaboration was sourced from Novartis clinical trials, and was approved by institutional review boards or ethics committees. All trials were conducted in accordance with the principles of the Declaration of Helsinki and the International Conference on Harmonisation guidelines for Good Clinical Practice.

## Data statement

Due to privacy requirements, we are unable to make the underlying data available for sharing.

